# Comparison among the first representative chloroplast genomes of *Orontium, Lasia, Zamioculcas*, and *Stylochaeton* of the plant family Araceae: inverted repeat dynamics are not linked to phylogenetic signaling

**DOI:** 10.1101/2020.04.07.029389

**Authors:** Abdullah, Claudia L. Henriquez, Furrukh Mehmood, Iram Shahzadi, Zain Ali, Mohammad Tahir Waheed, Thomas B. Croat, Peter Poczai, Ibrar Ahmed

## Abstract

The chloroplast genome provides insight into the evolution of plant species. We *de novo* assembled and annotated chloroplast genomes of the first representatives of four genera representing three subfamilies: *Lasia spinosa* (Lasioideae), *Stylochaeton bogneri, Zamioculcas zamiifolia* (Zamioculcadoideae), and *Orontium aquaticum* (Orontioideae), and performed comparative genomics using the plastomes. The size of the chloroplast genomes ranged from 163,770–169,982 bp. These genomes comprise 114 unique genes, including 80 protein-coding, 4 rRNA, and 30 tRNA genes. These genomes exhibited high similarities in codon usage, amino acid frequency, RNA editing sites, and microsatellites. The junctions JSB (IRb/SSC) and JSA (SSC/IRa) are highly variable, as is oligonucleotide repeats content among the genomes. The patterns of inverted repeats contraction and expansion were shown to be homoplasious and therefore unsuitable for phylogenetic analyses. Signatures of positive selection were shown for several genes in *S. bogneri*. This study is a valuable addition to the evolutionary history of chloroplast genome structure in Araceae.

## 1. Introduction

The chloroplast is an important double membrane bounded organelle that plays a crucial role in photosynthesis and in the synthesis of fatty acids and amino acids [1]. The chloroplast contains its own DNA, and replicates independently from the nuclear genome [1,2]. Chloroplast genomes mostly exhibit a quadripartite structure in which a pair of inverted repeats (IRa and IRb) separate large single copy (LSC) and small single copy (SSC) regions [1–4]. However, in some plant lineages, quadripartite structure is not observed due to the loss of one or two inverted repeats (IRs), for example in Taxodiaceae [5] and Fabaceae [6]. On the other hand, very short IRs are also reported, for example in Pinaceae [7]. Moreover, a mixture of linear and circular chloroplast genomes has also been observed [8].

The structure of chloroplast genomes is conserved regarding gene organization, gene content and intron content [1,9–12]. However, large scale events of gene rearrangement, gene loss/generation of pseudogenes, and intron loss are also reported in various plant lineages [10,13–17]. Inverted repeat contraction and expansion in chloroplast genomes create pseudogenes, cause gene duplication, or convert duplicate into single-copy genes [10,11]. Many other types of mutational events also take place within chloroplast genomes, including insertion-deletion (indels), substitutions, tandem repeat variations and variations in number and type of oligonucleotide repeats [11,18–20]. The single parent inheritance of the chloroplast genome – paternally in some gymnosperms and maternally in most angiosperms – along with adequate levels of polymorphism make it suitable for studies of evolution, domestication, phylogeography, population genetics, and phylogenetics [7,10,11,21–24].

Araceae is an ancient and large monocot plant family, comprising 114 genera and 3750 species [25]. This family is subdivided into eight subfamilies: Gymnostachydoideae, Orotioideae, Lemnoideae, Pothoideae, Monsteroideae, Lasioideae, Zamioculcadoideae, and Aroideae [22,26,27]. Advancements in next generation sequencing have made genomic resources available for species of Lemnoideae [28], Monsteroideae [18,29,30], and Aroideae [10,31,32]; chloroplast genomes of *Symplocarpus renifolius* Schott ex Tzvelev and *S. nipponicus* Makino of Orotioideae have also been reported [33,34]. Previous studies provide insight into some unique evolutionary events of chloroplast genomes in Araceae, including IR contraction and expansion, gene rearrangement, and positive selection [10,18,28]. Moreover, loss/pseudogenation of some important genes are also reported in the genus *Amorphophallus* Blume (Aroideae) [17]. Nevertheless, data for the chloroplast genome structure and evolution is still lacking for several major aroid clades, including the subfamilies Lasioideae and Zamioculcadoideae. Hence, new genomic resources are required to provide better insight into the evolution of chloroplast genomes in Araceae.

In the current study, we report *de novo* assembled and fully annotated chloroplast genomes of four species from three different subfamilies: *Lasia spinosa* (L.) Thwaites (Lasioideae), *Stylochaeton bogneri* Mayo and *Zamioculcas zamiifolia* (Lodd.) Engl. (Zamioculcadoideae), and *Orontium aquaticum* L. (Orontioideae). We performed comparative chloroplast genomics among these species. This study provides the first insights into the chloroplast genome structure of four genera of Araceae, along with the evolutionary rate of protein-coding genes and types of substitutions occurring within these chloroplast genomes.

## 2. Materials and Methods

### 2.1 Sample collection, DNA extraction and sequencing

We collected fresh and healthy leaves of four species (*L. spinosa, S. bogneri, Z. zamiifolia*, and *O. aquaticum*) from the Araceae Greenhouse at the Missouri Botanical Garden in St. Louis, Missouri. Whole genomic DNA was extracted from the collected leaves using Qiagen DNeasy Minikit (Qiagen, Germantown, Maryland, USA) with some modification following a previous approach [10,18]. DNA quality and quantity were confirmed by 1% agarose gel electrophoresis and Nanodrop (ThermoScientific, Delaware, USA). The libraries were constructed following manufacturer’s protocol of Illumina TruSeq kits (Illumina, Inc., San Diego, California) in the Pires laboratory at the University of Missouri, Columbia. The Illumina HiSeq 2000 platform was used to sequence qualified libraries from single end with 100 bp short reads at the University of Missouri DNA Core.

### 2.2 Genome assembly and annotation

The sequencing of these genomes generated 3.31 GB (*S. bogneri*) to 11.3 GB (*Z. zamiifolia*) of raw data (Table 1). The quality of the generated short read data was compared among species using FastQC and MultiQC [35,36]. The analyses confirmed high quality of the data with high average Phred score ranging from 35.69–37.6. The raw data of the sequenced four species were submitted to the Sequence Read Archive of the National Center for Biotechnology (NCBI) under SRA project number PRJNA613281. The generated sequence data were used to *de novo* assemble chloroplast genomes using Velvet v.1.2.10 [37] by generating contigs with various kmer values of 51, 61, 71, and 81, combined with the *de novo* assembly option of Geneious R8.1 [38] following previous studies [9,11,39]. The coverage depth analysis was performed by mapping the short reads to their respective *de novo* assembled chloroplast genomes by BWA mem [40]. The assembly of the genomes was then validated by visualizing in Tablet [41]. We observed issues at 4–5 points of repetitive regions, therefore, for further validation we used Fast-Plast v.1.2.2 following exactly the same procedure employed for the assembly of other Araceae species [10,18]. This helped us to corroborate the correct sequence at those points. The coverage depth analyses revealed that the average coverage depths of the genomes ranged from 92.7X–1021X. The *de novo* assembled chloroplast genomes were annotated by GeSeq [42], whereas tRNA genes were further verified by tRNAscan-SE v.2.0.3 [43] and ARAGORN v.1.2.38 [44] by selecting default parameters. The final annotated genomes were submitted to NCBI under specific accession numbers (Table 1). GB2sequin was used to generate five column tab-delimited files from the annotated genomes for NCBI submission [45]. The circular map of these genomes was drawn by OrganellarGenomeDRAW (OGDRAW) [46].

**Table 1.** Quality and quantity of whole genome short reads and coverage depth analyses of the *de novo* assembled genomes.

| Species | Data in GB | Whole genome reads | Phred Score | Plastome Reads | Average coverage | Maximum coverage | NCBI accession |
| --- | --- | --- | --- | --- | --- | --- | --- |
| *Orontium aquaticum* | 5.43 | 20,860,738 | 37.2 | 200,065 | 124.8 | 1347 | MT226773 |
| *Lasia spinosa* | 8.35 | 32,000,000 | 37.6 | 1,738,094 | 1021 | 1929 | MT226772 |
| *Zamioculcas zamiifolia* | 11.3 | 43,288,898 | 35.69 | 1,298,593 | 774.4 | 1293 | MT226775 |
| *Stylochaeton bogneri* | 3.31 | 12,709,376 | 37.39 | 148,896 | 92.7 | 749 | MT226774 |

### 2.3 Characterization, comparative analyses, and phylogenetic inference

We used Geneious R8.1 [38] to compare genomic features and determine amino acid frequency and codon usage. To visualize and compare the junctions of chloroplast genomes, we used IRscope with default parameters [47]. The integrated Mauve alignment [48] of Geneious R8.1 was used to analyze gene arrangement based on Colinear block analyses after removal of IRa from the genomes.

The Predictive RNA editors for Plant (PREP-CP) [49] program was used to determined RNA editing sites in the chloroplast genomes. We also analyzed microsatellites and oligonucleotide repeats using MISA (MIcroSAtellite) and REPuter, respectively. We determined microsatellites with repeat units as: mononucleotide repeats ≥ 10, dinucleotide ≥ 5, trinucleotide ≥ 4, tetranucleotide, pentanucleotide and hexanucleotide ≥ 3. The forward and reverse oligonucleotide repeats were determined with length ≥ 14 bp with 1 editing site, initially. Later, we removed all repeats that contained mismatches from the analyses and left only those repeat pairs that exhibited 100% similarities, following Abdullah et al. 2020 [50].

We determined transition substitutions (Ts), transversion substitutions (Tv) and their ratio (Ts/Tv) in 78 protein-coding genes. For this purpose, we concatenated protein-coding genes of all four species. The sequences of the concatenated protein-coding genes of *L. spinosa, S. bogneri*, and *Z. zamiifolia* were pairwise aligned to *O. aquaticum* by Multiple Alignment using Fast Fourier Transform (MAFFT). The substitution types were determined from each alignment in Geneious R8.1 [38].

We determined the rate of synonymous substitutions (K_s_), non-synonymous substitutions (K_a_) and their ratio (Ka/Ks) in 75 protein-coding genes. We extracted and aligned protein-coding genes from all four species. The chloroplast genome of *O. aquaticum* was used as a reference, and the rates of evolution of protein-coding genes were recorded. A similar approach was previously applied in other angiosperms [9,11,18,39,51]. The data were interpreted as: K_a_/K_s_ < 1, K_a_/K_s_ = 1, K_a_/K_s_ > 1, representing purifying, neutral and positive selection, respectively.

A phylogenetic analysis was performed among 30 species of Araceae, using *Acorus americanus* (Acoraceae) as an outgroup. MAFFT [52] on XSEDE v.7.402 in CIPRES [53] was used to align complete chloroplast genomes of all species after removal of one IR. The phylogeny was inferred based on this alignment after removal of indels using RAxML-HPC BlackBox v.8.2.12 [54] in CIPRES [53]. The details regarding the species that were used in the phylogenetic analysis are shown in table S1.

## 3. Results

### 3.1 Comparative genomics among *de novo* assembled chloroplast genomes

The sizes of the genomes ranged from 163,770 bp (*S. bogneri*) to 169,980 bp in *L. spinose*. The SSC region ranged from 13,967 bp (*O. aquaticum*) to 20,497 bp (*S. bogneri*); LSC ranged from 87,269 bp (*O. aquaticum*) to 91,357 bp (*Z. zamiifolia*); the size of each IR region ranged from 26,702 bp (*S. bogneri*) to 32,053 bp (*L. spinosa*) (Table 2). The chloroplast genomes of the four species were found to be highly conserved in terms of gene organization, gene content and intron content. This highly conserved structure was also confirmed using circular maps of the genomes (Figure 1), as well as from the Colinear Block analyses of Mauve (Figure 2). All species exhibited 114 unique genes, including 80 protein-coding, 30 tRNA, and 4 rRNA genes. We recorded 17 duplicated genes in the IRs of *S. bogneri* and *Z. zamiifolia*, and 18 duplicated genes in *O. aquaticum* and *L. spinosa*. In total, 18 intron-containing genes were observed, including 6 tRNA genes and 12 protein-coding genes. Among the intron-containing genes, 2 tRNA genes and 3 protein-coding genes are located in IRs. The size of introns showed some variation among species, whereas exons showed high similarity (Table S2). The *inf*A gene was found to be a pseudogene in all species. The GC content of the complete chloroplast genomes and of all regions showed high similarities among species, whereas fluctuation in GC content was observed within the different regions of the same chloroplast genome. The GC content of coding regions, rRNAs, and tRNAs also showed high similarities among species (Table 2).

**Table 2.**
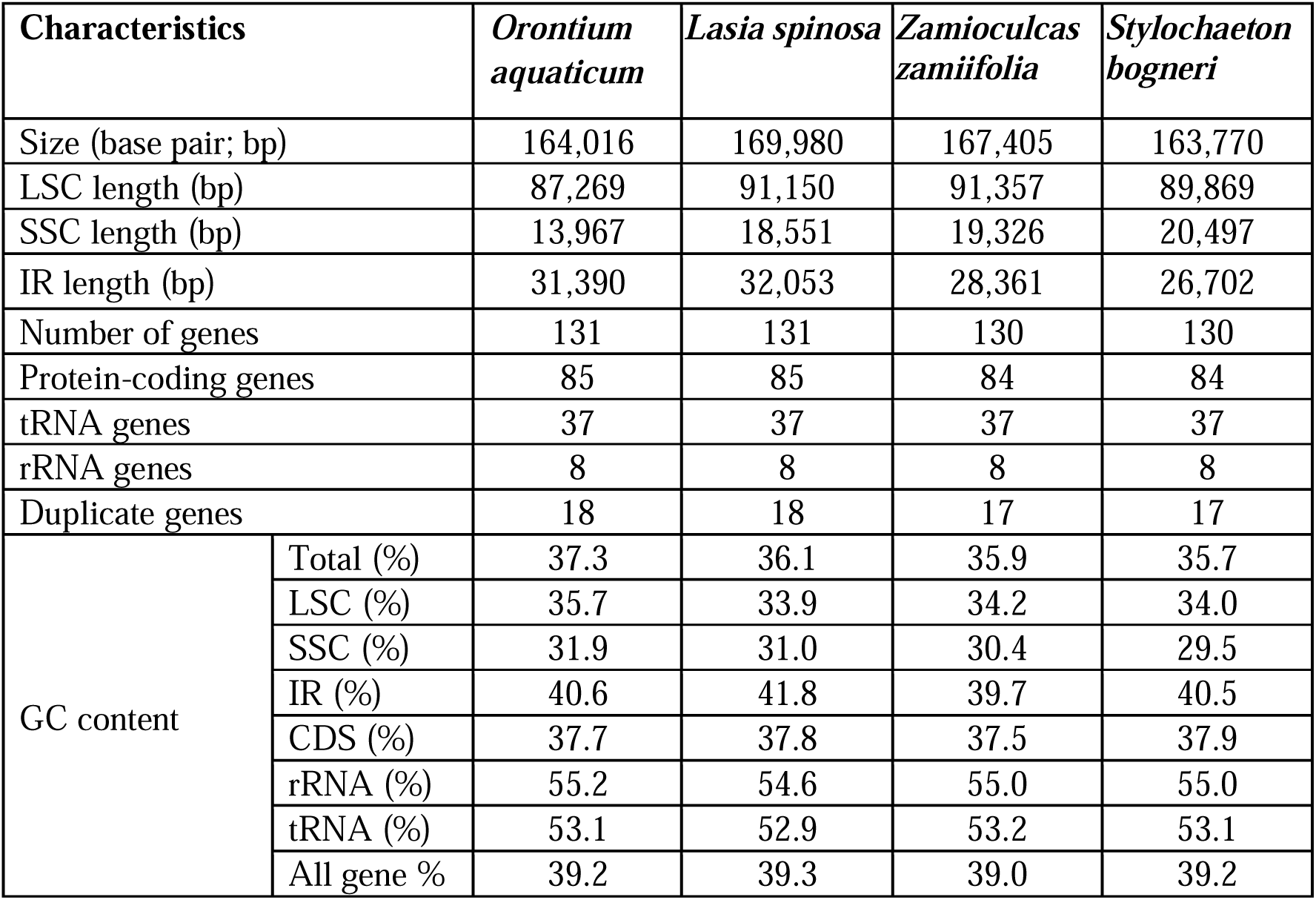
Genomic features of *de novo* assembled chloroplast genomes

**Figure 1.**
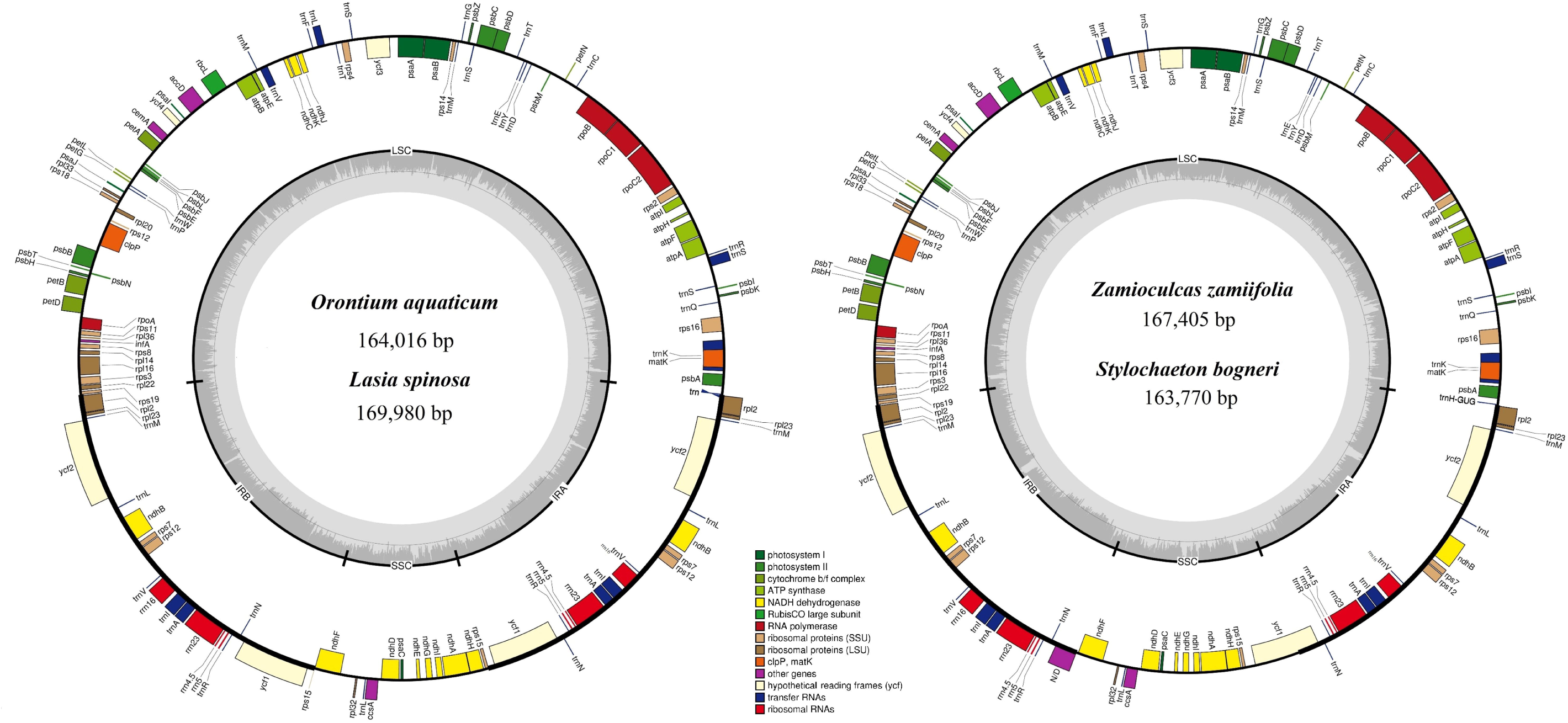
Circular maps of chloroplast genomes. Genes present inside the circle are transcribed counter clockwise, whereas genes present outside the circle are transcribed clockwise. Genes are colour coded based on functionality. LSC, IRb, SSC, and IRa of inner circle represent quadripartite structure of genomes.

**Figure 2.**
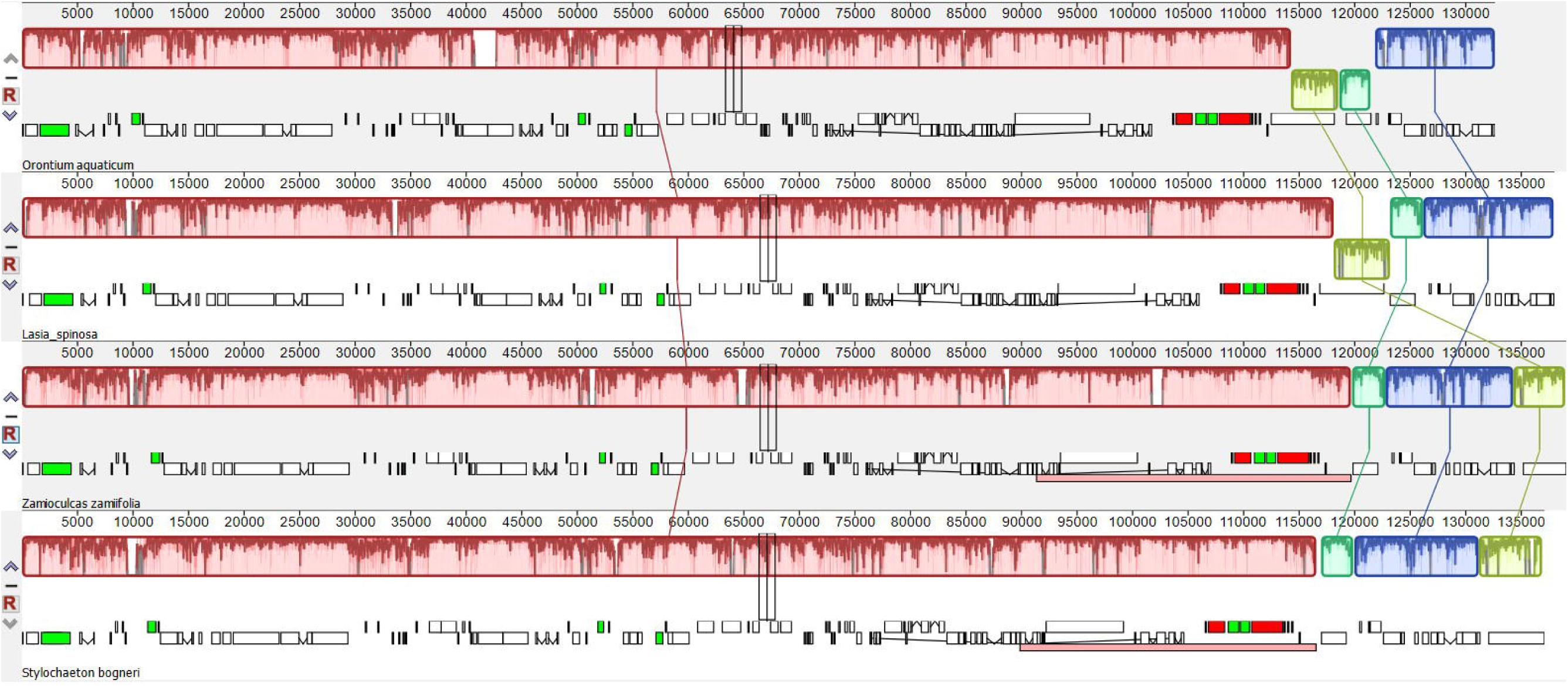
Colinear block-based analyses of gene arrangement in the chloroplast genomes. The black block: transfer RNA genes, green block: transfer RNA genes with introns, white block: coding-genes, red block: ribosomal RNA genes. Light green and dark green blocks show differential existence of *ycf1* and *rps15* due to inverted repeats contraction and expansion.

### 3.2 Inverted repeats contraction and expansion

The chloroplast genomes showed variations and similarities at the junctions of LSC/IR/SSC. The junctions of JLB (LSC/IRb) and JLA (IRa/LSC) showed similarities across all four species. The junctions of JSB and JSA were highly variable among species. The chloroplast genomes *O. aquaticum* and *L. spinosa* were found to be similar at these two junctions, and IR expansion led to duplication of the complete *ycf*1 gene and the origin of pseudogenes of *rps*15 at JSB. The chloroplast genomes of *S. bogneri* and *Z. zamiifolia* showed less expansion of IRs, which led to the origin of only a pseudogene of *ycf*1 at JSB. The integration of *ndh*F into IRb was only recorded in *L. spinosa*. At JLA in *O. aquaticum, trnH-GUG* was found to be completely in the LSC region 12 bp away from the junction, whereas other species showed integration of *trn*H*-*GUG into the IRa region from 6 bp to 11 bp. The complete details are presented in Figure 3.

**Figure 3.**
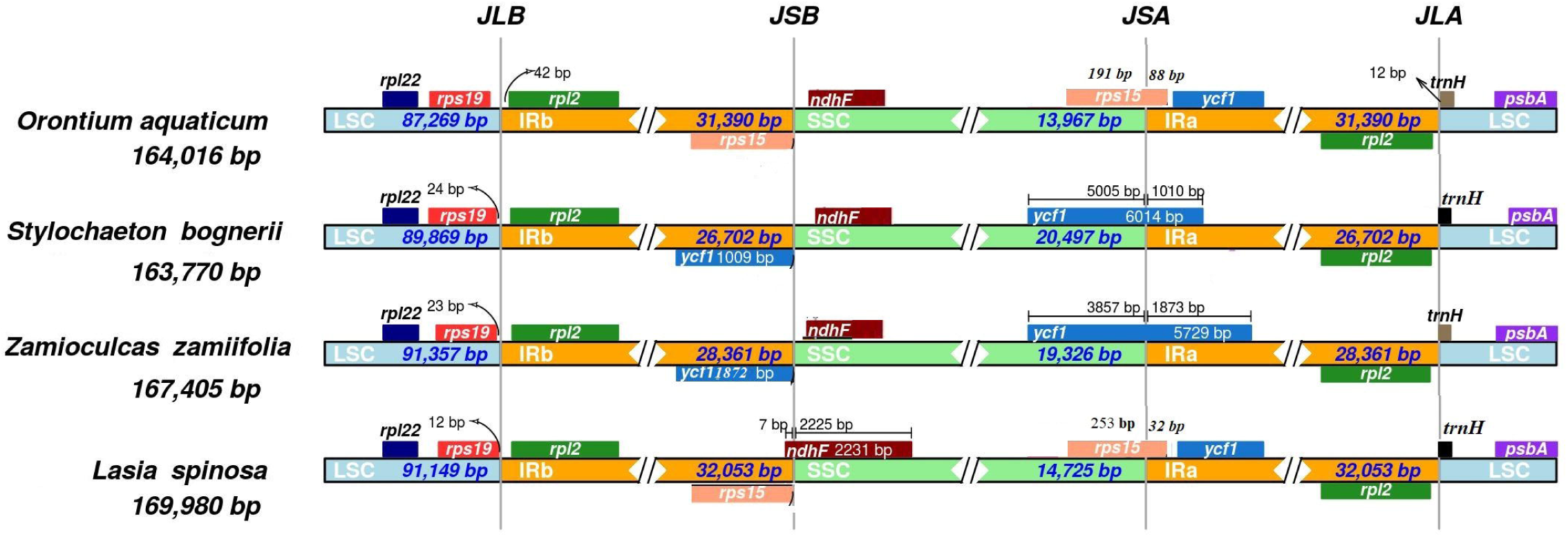
Comparison of quadripartite junction sites among chloroplast genomes of four assembled species. Genes present on top of track transcribe on the negative strand, whereas genes present below the track transcribe on the positive strand. The T scale bar shows integration of genes between two adjacent regions. The junctions of genomes are represented as: JLB: IRb/LSC, JSB: IRb/SSC, JSA: SSC/IRa, and JLA: IRa/LSC.

### 3.3 Analyses of codon usage, amino acid frequency and RNA editing

The codon usage analyses revealed high encoding efficacy for those codons that end with A/T as opposed to codons that end with C/G. We recorded a relative synonymous codon usage (RSCU) value ≥ 1 for most codons that end with A/T, whereas RSCU < 1 was recorded for codons that end with C/G (Table S3). The ATG codon is the most common start codon. However, we also observed ACG (in *rpl*2) and GTG (in *rps*19) as start codons. The amino acid frequency analyses revealed high encoding of leucine, whereas the rarest encoding was recorded for cysteine (Figure S1). The RNA editing analyses revealed the presence of 62–74 RNA editing sites in 19–21 genes (Table S4). The RNA editing sites were found in the same genes with a few exceptions: RNA editing sites were detected in *psa*B genes of only *Z. zamiifolia*, whereas the RNA editing site was not found in *rpl*20 of *S. bogneri* and *rps*8 of *L. spinosa*. Most of the RNA editing sites were recorded in *ndh*A, *ndh*B, *ndh*D, *rpo*A, *rpo*B, *rpo*C1, and *rpo*C2 (Table S4). ACG was found as a start codon in gene *rpl2* and the RNA editing analyses confirmed conversion of ACG codon to ATG. Most RNA editing sites were found to be related to conversion of serine to leucine. Moreover, almost all editing sites led to accumulation of hydrophobic amino acids in the polypeptide chain of proteins (Table S4).

### 3.4 Repeats analyses

The analyses of microsatellites revealed 104–146 repeats in the genomes. Most of the repeats existed in LSC followed by SSC and then IR (Fig. 4a). We recorded an abundance of mononucleotide repeats in all species, especially *Z. zamiifolia*. Dinucleotide repeats were in greater abundance in *O. aquaticum* and *L. spinosa*, whereas *S. bogneri* and *Z. zamiifolia* showed an abundance of mononucleotide repeats and tetranucleotide repeats. *Lasia spinosa, Z. zamiifolia* and *O. aquaticum* showed similarities in numbers of tri and tetranucleotide repeats in their respective genomes, but *S. bogneri* showed a lack of trinucleotide repeats relative to tetranucleotide repeats (Fig. 4b). Pentanucleotide and hexanucleotide repeats were in lower abundance than the other types of repeats and were completely lacking in Z. zamiifolia (Fig. 4a). Most repeats of all six microsatellite types were the A/T rather than G/C motif (Table S5). The analyses of oligonucleotide repeats revealed the existence of a higher number of forward and reverse repeats in all four species. We recorded most repeats in the LSC region as compared to the SSC and IR regions. We also found some shared repeats among the three regions of the chloroplast genomes (Figure 4c). The number of repeats ranged from 647 (*O. aquaticum*) to 1471 (*Z. zamiifolia*). We recorded high similarities in the numbers of forward and reverse repeats in *O. aquaticum* and *L. spinosa*, whereas in *S. bogneri* and *Z. zamiifolia* there was a higher abundance of forward repeats (Figure 4d). Most repeats ranged in length of 14–20 bp (Figure 4d), whereas the largest repeats varied from 39 bp (*L. spinosa*) to 75 bp (*S. bogneri*) (Figure 4e). The details about the position and number of repeats are provided in Table S6.

**Figure 4.**
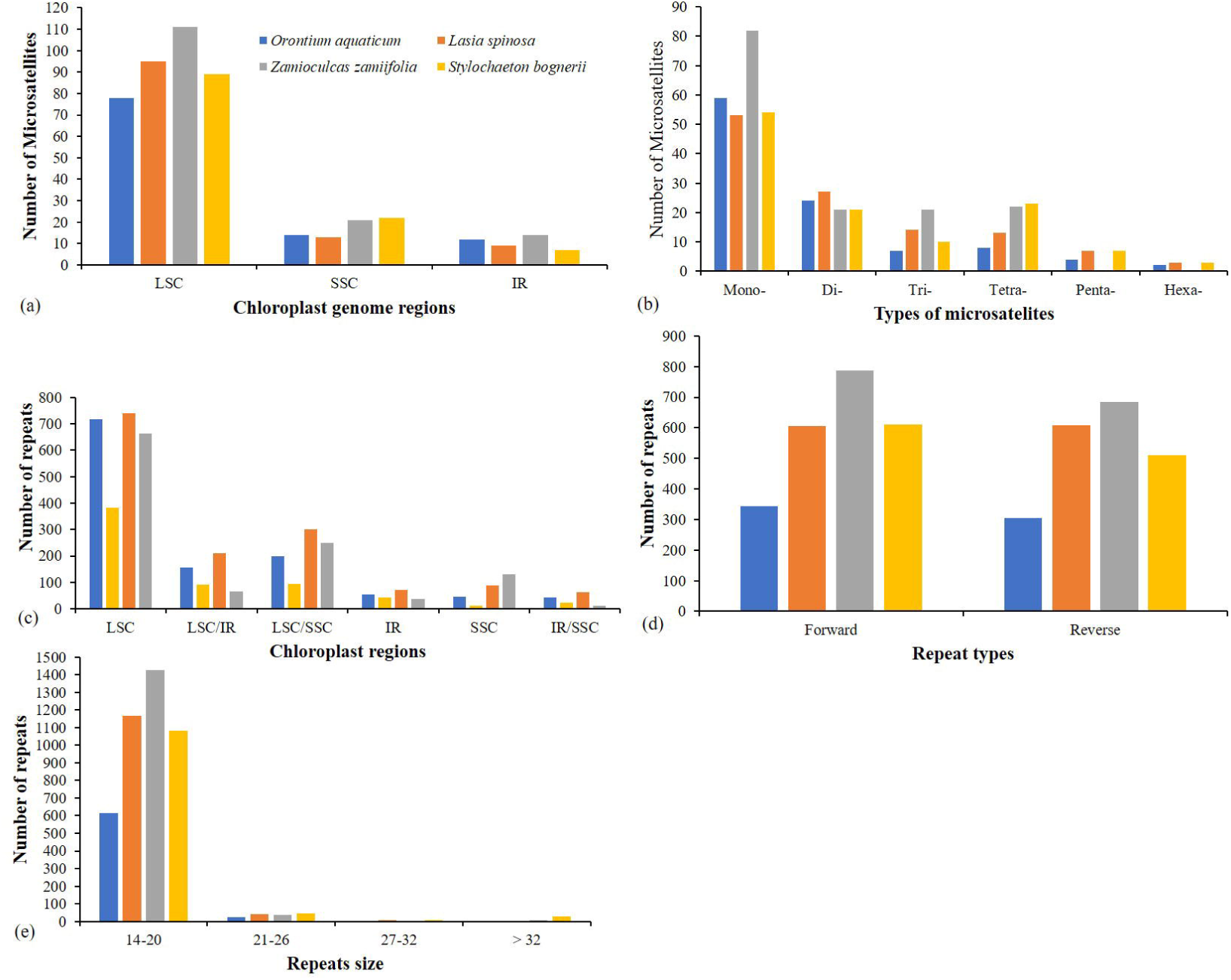
Comparison of repeats among chloroplast genomes of four species. (a) Microsatellites distribution in regions of chloroplast genomes. (b) Numbers of different types of microsatellites. (c) Distribution of oligonucleotide repeats in regions of chloroplast genomes. (d) Types of oligonucleotide repeats. (e) Number of repeats based on the size. LSC: large single copy, SSC: small single copy, IR: inverted repeats, LSC/SSC, LSC/IR, SSC/IR represent those repeats pairs in which one copy exists in one region and another copy in another region. 14-20, 21-26,27-32, >32 showed range of repeat size.

**Figure 5.**
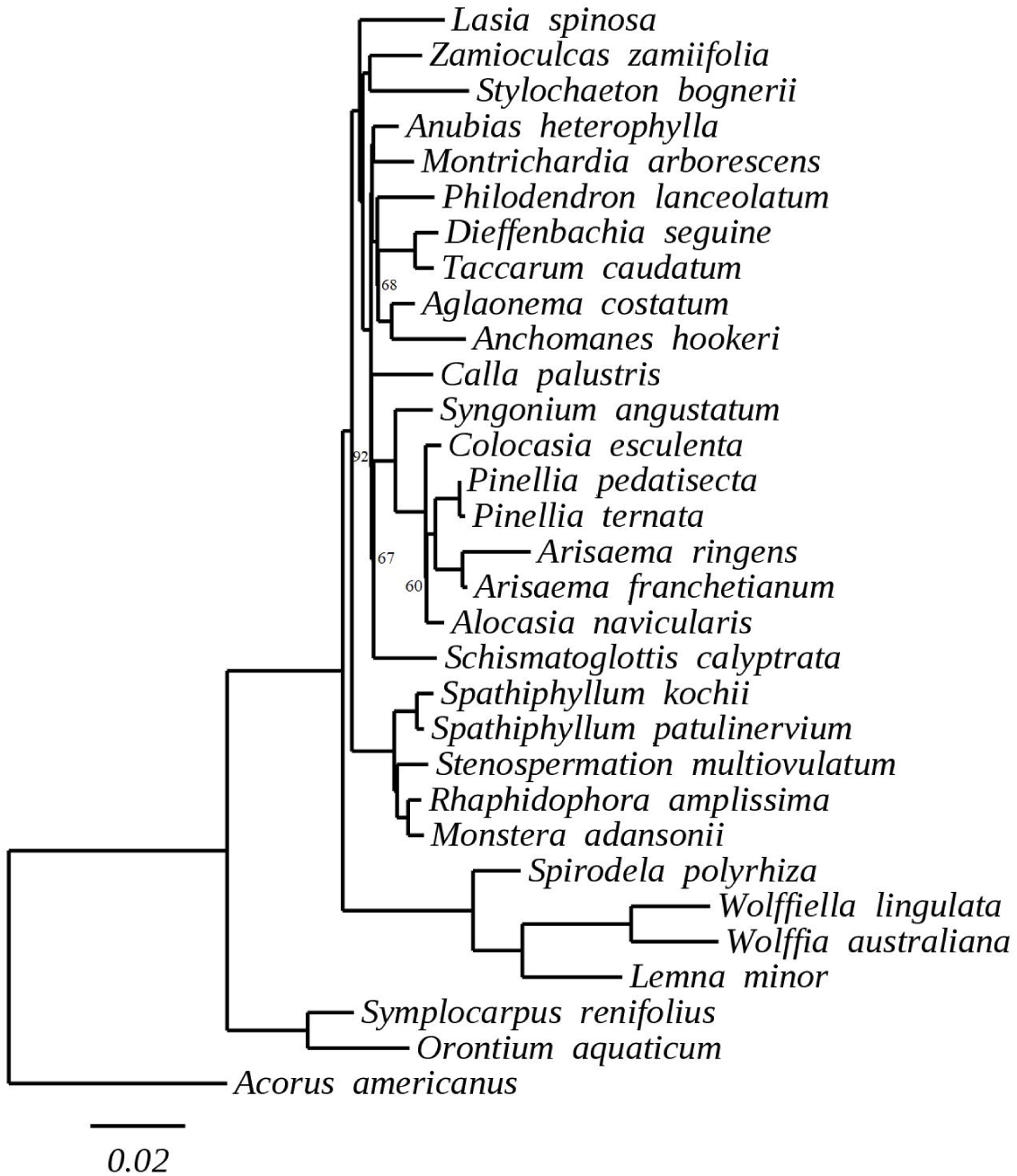
Maximum likelihood tree based on multiple alignment of 30 species of Araceae. We omitted bootstrapping support from those nodes which showed 100 bootstrapping support.

### 3.5 Analyses of substitutions types

We recorded a greater number of Ts than Tv substitutions. The ratio of Ts/Tv was 2.3, 2.03, and 2.15 in the genomes of *L. spinosa, S. bogneri*, and *Z. zamiifolia*, respectively. The majority of Ts were promoted by A/G rather than C/T, whereas the majority of Tv were found to be related to A/C and G/T rather than A/T and C/G (Table 3). For K_s_ and K_a_, we found a higher average of K_s_ than K_a_. Hence, on average, we recorded very low K_a_/K_s_ for all genes, which shows that purifying selection is acting on these genes. The average values recorded for the different groups of genes were: photosystem I group (K_s_ = 0.1677, K_a_ = 0.0125, and K_a_/K_s_ = 0.1211), photosystem II (K_s_ = 0.1208, K_a_ = 0.0085, and K_a_/K_s_ = 0.0671), cytochrome group (K_s_ = 0.1757, K_a_ = 0.0298, and K_a_/K_s_ = 0.2012), ATP synthase group (K_s_ = 0.1466, K_a_ = 0.0188, and K_a_/K_s_ = 0.1337), ribosomal small subunit group (K_s_ = 0.1620, K_a_ = 0.0633, and K_a_/K_s_ = 0.3491), ribosomal large subunit (K_s_ = 0.1639, K_a_ = 0.0677, and K_a_/K_s_ = 0.4612), NADPH dehydrogenase group (K_s_ = 0.1711, K_a_ = 0.0781, and K_a_/K_s_ = 0.3935), and RNA polymerase group (K_s_ = 0.1813, K_a_ = 0.3228, K_a_/K_s_ = 0.1800) (Table S7). Some genes showed neutral selection (K_a_/K_s_ = 1) in all species, including *ndh*K, *pet*L, *rpl*16, *ndh*F, *ndh*H, and *rps*15. Interestingly, we found evidence for positive selection in three genes (*ycf*2, *clp*P, *rpl*36) in only *S. bogneri* (Table S7).

**Table 3.** Transition and transversion substitutions in protein-coding genes.

| **Substitution types** | *Lasia spinosa* | *Zamioculcas zamiifolia* | *Stylochaeton bogneri* |
| --- | --- | --- | --- |
| A/C | 478 | 466 | 560 |
| C/T | 1375 | 1267 | 1457 |
| A/G | 1429 | 1316 | 1540 |
| A/T | 274 | 254 | 315 |
| C/G | 160 | 156 | 193 |
| G/T | 305 | 322 | 406 |
| Ts/Tv | 2.3 | 2.15 | 2.03 |

### 3.6 Phylogenetic inference of Araceae

A phylogenetic tree was reconstructed using 31 species based on an alignment of 93,794 nucleotide sites, of which 13,488 were found to be parsimony informative; 8,843 showed distinct site pattern while the remaining sites (67,229 sites) were shared in all species. The resulting phylogeny shows the monophyly of Lasioideae, Zamioculcadoideae and Orontioideae, with Zamioculcadoideae forming a clade with *Stylochaeton*.

## Discussion

In the current study, we report *de novo* assembled and fully annotated chloroplast genomes of four species from three subfamilies of Araceae. Comparative chloroplast genomics revealed high similarities in gene content across all species. However, the sizes of these genomes varied due to the variable length of intergenic spacer regions (IGS) and IRs contraction and expansion. The substitution analyses revealed Ts > Tv and K_s_ > K_a_. The phylogenetic analysis confirmed the monophyly of Lasioideae, Orontioideae and Zamioculcadoideae.

The chloroplast genomes of the four species showed a highly conserved structure in terms of gene content, intron content and gene organization. Similar gene contents were also reported in other subfamilies of Araceae [10,18,28,34]. Moreover, highly conserved chloroplast genomes are reported in other angiosperms [10,18,55]. However, loss of some important protein-coding genes and tRNA genes has been reported in the genus *Amorphophallus* (Aroideae, Araceae), which might be specific to this genus. The *inf*A gene encodes translation initiation factor I, but we found this gene to be non-functional in all species. This gene is also reported to be non-functional or absent in the chloroplast genomes of other angiosperms including species of Araceae [9–11,18,28,55]. Hence, it is suggested that this gene is either transferred to the nuclear genome as an active functional gene, or a functional copy of this important gene might already exist in the nuclear genome [31,56]. We observed duplication of *ycf*1 genes or origination of pseudogenes of *ycf*1 and *rps*15 due to IR contraction and expansion. The duplication of *ycf*1 and/or *rps*15 is also reported in species of the subfamily Lemnoideae [28] and two species (*Anchomanes hookeri* Schott. and *Zantedeschia aethiopica* Spreng.) of Aroideae [10].

Together with previously published aroid chloroplast genomes, the chloroplast genomes of the current study reveal that IR contraction and expansion might be a species-specific event, opposed to synapomorphies of subfamilies. The duplication of *ycf*1 and origination of the *rps*15 pseudogene in *O. aquaticum* is not observed in species of *Symplocarpus* Salisb. ex Nutt. [34], and both genes are found in the SSC region. The complete duplication of *ycf*1 and *rps*15 is observed in Lemnoideae species [28]; in *S. bogneri* and *Z. zamiifolia*, duplication of partial *ycf*1 is observed, and *rps*15 completely exists in the SSC region. Moreover, complete duplication of *ycf*1 and partial duplication of *rps*15 was observed in *L. spinosa*, similar to *O. aquaticum*. This data suggests that IR contraction and expansion is highly flexible over evolutionary time, and that similar IR boundary architecture across lineages can be the result of homoplasy. In other angiosperms, differential IR contraction and expansion has also been seen in species of the same genus such as *Aquilaria* Lam. [57,58]. This observation is contradictory to a previous study in which resemblance of IR junctions was suggested for phylogenetic inference [59]. However, these authors did not provide conclusive results and suggested further study in wide range of germplasms.

The high RSCU value has also been previously reported for those codons that contain A/T at 3′ end instead of C/G, which might be due to the high AT content of the chloroplast genomes [3,11,18,55]. We found an abundance of leucine and isoleucine, whereas cysteine was found to be the rarest amino acid. These results are in agreement with previous studies of angiosperms, including the family Araceae [3,9,16,18,28]. We found high similarities in RNA editing sites. Most of the conversions by RNA editing led to the addition of hydrophobic amino acids to polypeptide chains of proteins. A similar pattern of conversion has been noted in the chloroplast genomes of other angiosperms [11,16,19]. We found ACG as a start codon for *rpl*2 instead of ATG, and RNA editing analyses confirmed the conversion of ACG to ATG. This is in agreement with a previous study [60].

Oligonucleotide repeats generate mutations in genomes [11,31,50,61]. Hence, these repeats are suggested to be used as a proxy to identify mutational hotspots [11,31,50]. In the current study, we analyzed forward and reverse repeats, as these repeats are considered important for the generation of mutations [11,31,50] and show up to 90% co-occurrence with substitutions as reported in the plant family Malvaceae [50]. Previously, high similarities were reported in repeat numbers and types in some angiosperms [3,4,9,11,19,55]. However, previous studies also show significantly variable number of repeats in other species of Araceae, which does not correlate with genome size or phylogenetic position [10,18]. Here, our result agrees with previous studies of Araceae. Notably, penta- and hexanucleotide repeats were completely absent in *Z. zamiifolia*, while the number of mononucleotide repeats was almost twice as high as in the other genera. In the current study, we changed the criteria for repeat determination following a recent study of Abdullah et al. 2020 [50], and identified repeats of exact match ≥ 14. However, our result still agrees with a previous study in that repeats in the chloroplast genomes of Araceae are independent of genome size and phylogenetic position.

We observed a greater amount of Ts than Tv substitutions within the protein-coding genes, as expected in DNA sequences [62]. However, fewer Ts than Tv has also been reported previously in chloroplast genomes [9,39,63]. Higher Ts than Tv might occur due to genome composition and codon characteristics [64]. Moreover, higher Ts than Tv was also reported in the complete chloroplast genome of *Dioscorea polystachya* Turcz. [65] and in the coding sequences of the species of Lemnoideae (Araceae). This suggests that species of the plant family Araceae are consistent regarding the existence of higher Ts as compared to Tv.

We observed K_a_/K_s_ < 1 due to higher K_s_ than K_a_ for most of the protein-coding genes. These results are consistent with previous studies of angiosperm chloroplast genomes, including the family Araceae, as purifying selection pressure mostly acts on the genes of chloroplast genomes [9,11,18,55]. However, a higher K_a_/K_s_ was also reported in some species of Araceae in which most of the genes were under positive selection [34]. We also found three genes under positive selection in the chloroplast genome of *S. bogneri*, including *ycf*2, *clp*P, and *rpl*36, which might be due to the different types of stresses faced by these species in their respective ecological niches. These genes were also found to be under positive selection in various other species [11,16,55,66,67].

This study presents a comparison of the complete chloroplast genomes of four species representing three subfamilies of Araceae including Orontioideae, Lasioideae and Zamioculcadoideae, and the independent taxon *S. bogneri*. IR contraction and expansion appears to be homoplasious among these taxa, precluding the use of the IR architecture in phylogenetic analyses. Unique features of members of the *Stylochaeton* clade [26], represented here by *Z. zamiifolia* (Zamioculcadoideae) and *Stylochaeton* Lepr., include repeat types and amounts in both species, and signatures of positive selection in several genes in *Stylochaeton*.

## Authors Contribution

Sample collection, DNA extraction and sequencing: CH and TC; Genomes assembly, coverage depth analyses and annotations: A, CH, and ZA; Data analyses: A, FM and IS; Data interpretation: A and FM; Conceptualization: A, PP, IA, MTW; Data curation: A, CH, FM; Project administration: A, CH; Writing – Original Draft: A; Writing - Review & Editing: CH, IA, PP, MTW; Supervision: PP and IA

## Acknowledgment

Authors are thankful for funding and laboratory support to Dr. Barbara Schaal at Washington University in St. Louis and Dr. J. Chris Pires at the University of Columbia, Missouri. Authors are also thankful to Dr. Tatiana Arias for valuable help in the laboratory and data processing. In the aroid greenhouse at the Missouri Botanical Garden, Emily Colletti provide help with living material and authors like to thank for it.

## Figure Legend

**Figure S1**. Comparison of amino acid frequency among four species of Araceae.

