## Supplementary figures and images for "Comparison among the first representative chloroplast genomes of *Orontium, Lasia, Zamioculcas*, and *Stylochaeton* of the plant family Araceae: inverted repeat dynamics are not linked to phylogenetic signaling"

### Supplementary Figure 1

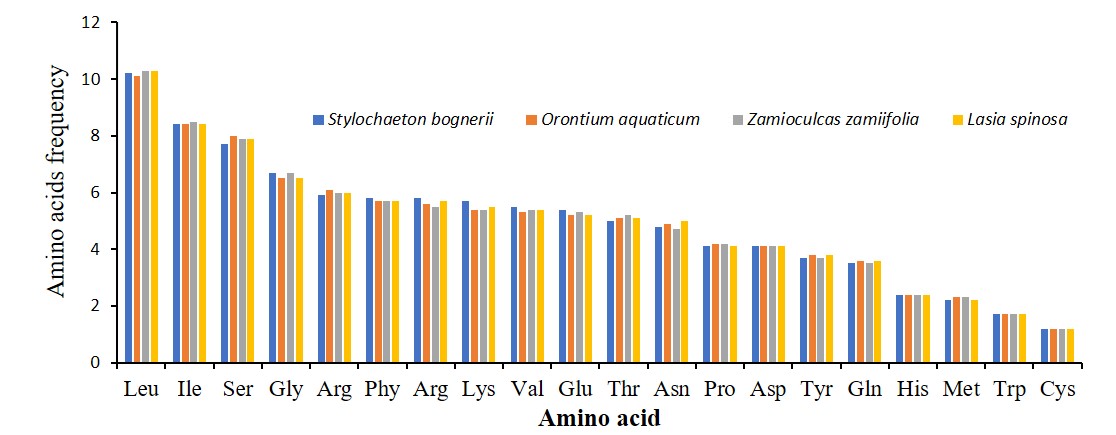
