## Supplementary Table 1 for "Comparison among the first representative chloroplast genomes of *Orontium, Lasia, Zamioculcas*, and *Stylochaeton* of the plant family Araceae: inverted repeat dynamics are not linked to phylogenetic signaling"

Table S1. GenBank accession number of the species used in phylogenetic inference

| S. No | Species | Accession |
| --- | --- | --- |
| 1 | *Acorus americanus (outgroup)* | EU273602 |
| 2 | *Colocasia esculenta* | JN105689 |
| 3 | *Lemna minor* | DQ400350 |
| 4 | *Spirodela polyrhiza* | JN160603 |
| 5 | *Wolffiella lingulata* | JN160604 |
| 6 | *Wolffia australiana* | JN160605 |
| 7 | *Spathiphyllum kochii* | KR270822 |
| 8 | *Pinellia ternata* | KR270823 |
| 9 | *Symplocarpus renifolius* | KY039276 |
| 10 | *Arisaema ringens* | MK111107 |
| 11 | *Dieffenbachia seguine* | NC_027272 |
| 12 | *Anubias heterophylla* | MN046884 |
| 13 | *Arisaema franchetianum* | MN046885 |
| 14 | *Schismatoglottis calyptrata* | MN046892 |
| 15 | *Pinellia pedatisecta* | MN046890 |
| 16 | *Philodendron lanceolatum* | MN551187 |
| 17 | *Taccarum caudatum* | MN046895 |
| 18 | *Montrichardia arborescens* | MN046889 |
| 19 | *Aglaonema costatum* | MN046881 |
| 20 | *Alocasia navicularis* | MN046882 |
| 21 | *Syngonium angustatum* | MN046894 |
| 22 | *Anchomanes hookeri* | MN551188 |
| 23 | *Calla palustris* | MN046887 |
| 24 | *Monstera adansonii* | MN046888 |
| 25 | *Rhaphidophora amplissima* | MN046885 |
| 26 | *Stenospermation multiovulatum* | MN046893 |
| 27 | *Spathiphyllum patulinervum* | MN046890 |
