## Supplementary Table 2 for "Comparison among the first representative chloroplast genomes of *Orontium, Lasia, Zamioculcas*, and *Stylochaeton* of the plant family Araceae: inverted repeat dynamics are not linked to phylogenetic signaling"

**Table S2. The lengths of introns and exons in the intron containing genes of de novo assembled species**

***Lasia spinosa***

______________________________________________________________________________

Gene	Strand	Start	End	Exon I	Intron I	Exon II	Intron II	Exon III

______________________________________________________________________________

*trnK-UUU*	-	1990	4625	37	2563	36

*rps16*	-	5338	6617	42	1043	237

*trnS-CGA*	+	10923	11692	31	709	60

*atpF*	-	13657	15152	145	950	401

*rpoC1*	-	22858	25649	423	740	1629

*ycf3*	-	46574	48637	126	751	226	806	155

*trnL-UAA*	+	52052	52665	35	529	50

*trnV-UAC*	-	57196	57866	39	579	37

*clpP*	-	76300	78379	71	836	292	638	243

*petB*	+	81336	82783	6	800	642

*petD*	+	83066	84292	8	744	475

*rpl16*	-	87923	89496	9	1166	399

*rpl2*	-	91204	92685	391	663	431

*ndhB*	-	102157	104358	771	675	756

*trnE-GAU*	+	109987	110998	32	940	40

*trnA-UGC*	+	111063	111934	38	799	35

*ndhA*	-	134199	136379	553	1089	539

*trnA-UGC*	-	149196	150067	37	799	36

*trnE-GAU*	-	150132	151143	32	940	40

*ndhB*	+	156772	158973	771	675	756

*rpl2*	+	168442	169926	391	663	431

***Orontium aquaticum***

_____________________________________________________________________________

Gene	Strand	Start	End	ExonI	IntronI	ExonII	IntronII	ExonIII

_____________________________________________________________________________

*trnK-UUU*	-	1642	4198	38	2520	36

*rps16*	-	4851	6403	42	1324	228

*trnS-CGA*	+	9907	10648	31	710	60

*atpF*	-	12660	14028	145	814	410

*rpoC1*	-	21848	24656	423	757	1629

*ycf3*	-	44842	46857	126	760	226	749	155

*trnL-UAA*	+	50107	50786	35	595	50

*trnV-UAC*	-	54298	54973	33	584	59

*clpP*	-	72743	74828	71	809	292	671	243

*petB*	+	77782	79231	6	802	642

*petD*	+	79434	80662	8	744	475

*rpl16*	-	85607	84102	9	1098	399

*rpl2*	-	87312	88802	391	669	431

*ndhB*	-	97835	100067	777	700	756

*trnE-GAU*	+	105697	106720	32	952	40

*trnA-UGC*	+	106785	107656	37	799	36

*ndhA*	-	128910	131149	553	1148	539

*trnA-UGC*	-	143630	144501	37	799	36

*trnE-GAU*	-	144566	145589	32	952	40

*ndhB*	+	151219	153451	777	700	756

*rpl2*	+	162484	163974	391	669	431

***Stylochaeton* *bogneri***

___________________________________________________________________________

Gene	Strand	Start	End	Exon I	IntronI	ExonII	IntronII	ExonIII

_____________________________________________________________________________

*trnK-UUU*	-	1750	4388	37	2567	35

*rps16*	-	5206	6507	42	1065	195

*trnS-CGA*	+	11330	12126	31	706	60

*atpF*	-	14079	15447	145	823	401

*rpoC1*	-	23322	26104	423	731	1629

*ycf3*	-	46272	48299	126	755	226	766	155

*trnL-UAA*	+	51856	52450	35	510	50

*trnV-UAC*	-	57084	57748	33	575	57

*clpP*	-	76709	77305	71	809	292	671	231

*petB*	+	80140	81614	6	802	642

*petD*	+	81815	88197	8	744	475

*rpl16*	-	86499	84102	9	1098	399

*rpl2*	-	89919	91404	391	664	431

*ndhB*	-	100740	102947	777	675	756

*trnE-GAU*	+	108625	109641	32	945	40

*trnA-UGC*	+	109706	110581	37	803	36

*ndhA*	-	127518	129721	556	1109	539

*trnA-UGC*	-	143059	143934	37	803	36

*trnE-GAU*	-	143999	145015	32	945	40

*ndhB*	+	150693	152900	777	675	756

*rpl2*	+	162236	163721	391	664	431

***Zamioculcas zamiifolia***

______________________________________________________________________________

Gene	Strand	Start	End	ExonI	IntronI	ExonII	IntronII	ExonIII

______________________________________________________________________________

*trnK-UUU*	-	1830	4471	38	2568	36

*rps16*	-	5258	6572	42	1079	195

*trnS-CGA*	+	11667	12461	31	704	60

*atpF*	-	14417	15777	145	815	401

*rpoC1*	-	23434	26216	423	731	1629

*ycf3*	-	46092	48122	126	759	226	765	155

*trnL-UAA*	+	52011	52631	35	536	50

*trnV-UAC*	-	56682	57347	39	592	35

*clpP*	-	76327	78418	71	827	292	659	243

*petB*	+	81297	82740	6	802	642

*petD*	+	82942	84194	8	744	475

*rpl16*	-	87775	89712	9	1098	399

*rpl2*	-	91413	92898	391	664	431

*ndhB*	-	103176	105385	777	677	756

*trnE-GAU*	+	110980	111990	32	939	40

*trnA-UGC*	+	112055	112932	37	805	36

*ndhA*	-	130541	132791	553	1159	539

*trnA-UGC*	-	145831	146708	37	805	36

*trnE-GAU*	-	146773	147783	32	939	40

*ndhB*	+	153378	155587	777	677	756

*rpl2*	+	165865	167350	391	664	431
